# Cell fate determination by Lamarckian molecule-inheritance and chance

**DOI:** 10.1101/618199

**Authors:** Frank J. Bruggeman, Jaap Schouten, Daan H. de Groot, Robert Planqué

## Abstract

Single, isogenic cells can differ in their survival and adaptation capacity. This phenotypic diversity is generally due to stochastic molecular events. Since mother cells on average pass half of their molecular content on to their daughters, the states of progeny cells strongly correlate with that of mother cells (Lamarckian inheritance). Why a particular cell deviates qualitatively from others therefore requires consideration of chance events along its ancestral lineage. Here we develop theory to understand cellular heterogeneity in terms of stochastic ancestral events of molecule synthesis, molecule degradation and cell divisions. We find that cell growth stochasticity has profound consequences for molecular heterogeneity in isogenic populations of cells, especially for long-lived molecules such as proteins. For instance, the lower bound on noise in molecule copy numbers that has often been observed experimentally is shown to be solely determined by the probability distribution for the generation times of cells. Thus, copy-number noise is unavoidable, even in high-copy number circuits. Stochastic cell-fate and cell-differentiation decisions are therefore not necessarily due to noise in genetic circuits. We conclude that consideration of past chance events along cell lineages improves our understanding of how adaptive and maladaptive phenotypic heterogeneity arises in populations of isogenic cells.

## 1 Introduction

Random fluctuations in molecule concentrations may cause a particular cell to deviate in behaviour relative to others, even its close ancestors (Rosenfeld et al., 2005). Such phenotypic heterogeneity emerges spontaneously in isogenic cell populations (Paulsson, 2004). It may lead to the formation of subpopulations of cells, which in some cases spontaneously switch between phenotypic states at random times (Norman et al., 2015).

Whether fluctuations in a specific kind of molecule can have a significant phenotypic influence depends, amongst other factors, on its life time (Choi et al., 2008). Whereas some molecules live for a short period, because of rapid degradation, others exist for a long time and can only effectively reduce in concentration due to cell division events. For such stable molecular species, some copies in a cell were made by this cell itself, while others were inherited from its mother, grandmother, etc. The transition of a cell to another state, due to a deviating molecular content, can therefore be caused by events in its ancestral cells. For a stable molecule, at the end of a cell cycle, on average 50% of the cell’s molecules came from its ancestors. If an ancestral cell had a high molecular content because of a stochastic event, such as a higher overall synthesis rate or a longer life time, then this causes the daughter cell to be born with high molecule counts as well. In short, past chance events can have phenotypic consequences for extant cells, a true case of Lamarckian inheritance.

This phenomenon has been illustrated for the *lac* operon in *E. coli* (Choi et al., 2008). When *E. coli* grows in an environment with glucose and lactose, it prefers glucose and grows on lactose only when glucose has been depleted. Since *E. coli* requires a lactose permease to import lactose and lacks a lactose sensor, cells only initiate growth on lactose if a lactose permease was already produced purely by chance during growth on glucose, or after glucose depleted. This mechanism might be functional for all nutrients that cannot pass the membrane freely by diffusion and for which cells lack a sensing mechanism (van Boxtel et al., 2017).

Another example of the importance of past chance events is the occurrence of phenotypic diversification in isogenic cell populations. Phenotypic diversification can be reversible or irreversible. Persister formation in *E. coli* is an example of the reversible case, in which cells randomly switch between a growing and a hardly-growing antibiotic-tolerant phenotype (Norman et al., 2015). If this population is confronted with a sudden extinction-threatening condition, the persister cells have a higher chance of survival. Such behaviour can result for instance from the stochastic dynamics of toxin/antitoxin systems and fluctuations in the alarmone molecule ppGpp (Maisonneuve et al., 2013). In the irreversible case, spontaneous, random switching between phenotypes does not occur. During growth-supporting conditions one phenotype, e.g., a stress-resistant spore (Levine et al., 2012), is made via a chance event. The spore can survive harsh conditions, during which the growing phenotype dies, and can resuscitate again when conditions improve.

Here we present a theory that describes how the variability in molecular content of a single cell relates to random molecular events in itself and in any of its ancestors. We consider a description of the stochasticity of molecular circuits, cell growth and division; both of these influence the stochasticity of molecule copy numbers. At the level of molecular circuits, random fluctuations arise because of asynchronous molecule synthesis and degradation events that are influenced by fluctuating regulators and feedback circuitry. Copy-number stochasticity also arises because some cells have longer generation times than others, causing them to have more molecules at the end of their lifetime. In addition, random partitioning of molecules over two daughter cells during cell division, causes variation in molecule number between sisters. In order to obtain an ancestral perspective on molecule copy number in a single cell, we have to consider for each of its molecules the probability that they were made a certain number of generations ago, have not been degraded since, and ended up in the focal cell after each cell division. If we would achieve this, we would have a complete understanding of molecular fluctuations in living cells.

The outline of this paper is as follows. We start by illustrating how phenotypic heterogeneity can arise by considering the molecular stochasticity in a single cell that is part of a population of cells in balanced growth. Next, we express the probability that a single cell contains a certain number of molecules in terms of molecular chance events in any of its ancestral cells. We derive a loss rate of molecular memory in terms of molecular and cellular parameters. Then we apply the theory to stable molecules, for which ancestral effects are most important. We characterise the ancestral contribution to the statistics of molecule copy numbers of a single cell for different simple molecular circuits for protein synthesis. In all these cases, we find a non-negligible contribution of the stochasticity of cell growth and division events on the molecular stochasticity of a single cell. The randomness in generation times of cell (i.e., cell cycle durations) causes macroscopic, deterministic models, and the variance estimates deriving from those, to be inexact. We discuss how significant those deviations are for making predictions. Another finding is that molecules display stochasticity that is entirely due to cellular, rather than molecular, chance events. This explains the existence of a lower noise floor that is often observed experimentally. Since the noise floor turns out to be due to cellular stochasticity, it has consequences even for abundant proteins, such as those involved in metabolic networks. Cell-fate decisions are therefore not limited to low-copy number genetic circuits, but can also arise in such high-copy number circuits.

## 2 Results

### 2.1 Cell fate decisions

Phenotypic diversification of a population of isogenic cells, which we study in this paper, is explained in Figure 1A. Imagine a cell that can exist in one of two phenotypes. We shall assume that a cell’s fate, i.e., the phenotype it displays, depends on whether a particular molecule *Y,* such as a transcription factor, is above or below a critical concentration (Figure 1A). The level *y* of *Y* is assumed to be entirely set by the level *x* of another regulating molecule *X*, so *y* = *f* (*x*) for some suitable function *f*. Since fluctuations occur in *X*, of which some will propagate to *Y,* the noise in *Y* is partially due to noise in *X*. High noise in *X* might then cause some cells to be in phenotypic state 1 and others in state 2 (Figure 1B). To understand this diversification, we have to understand why cells have deviating levels of *X*. For proteins, which are generally long-lived, the copy number of molecules found in a cell depends on past molecular and cellular events. Hence, we have to look back in time along the cell lineage to find the ancestral cells that had large enough fluctuations of *X* (Figure 1C). An explicit experimental example of this phenomenon can be found in Choi et al. (2008).

**Figure 1:**
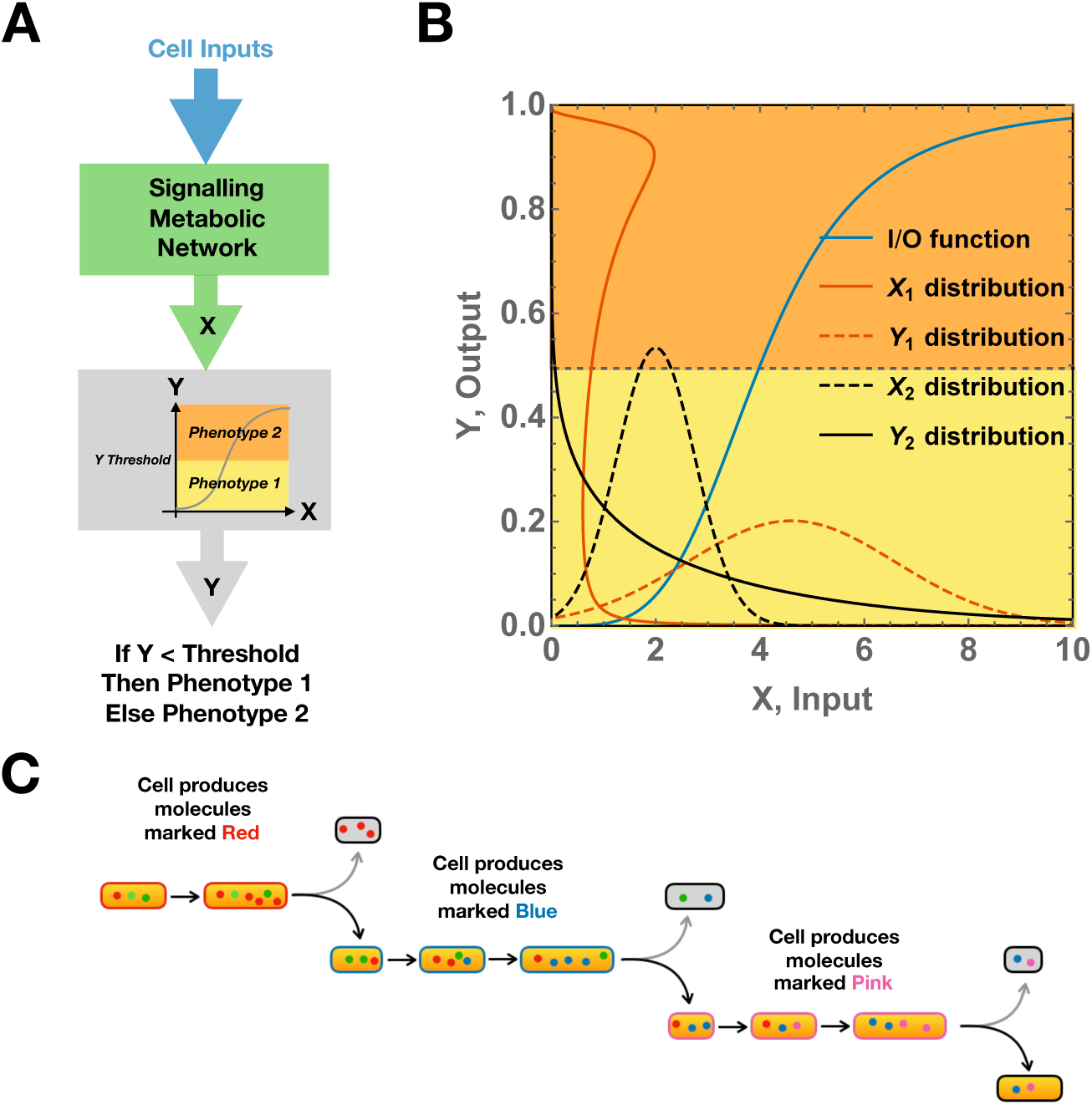
How variable molecular states can affect a cell’s fate. **A.** A schematic overview of the cellular decision making process of a cell. The regulatory molecule *X* decides on a cell’s fate, i.e., phenotype 1 or 2, by inducing changes that either cause *Y* to exist above or below a threshold level for two phenotypic states. **B.** Fluctuations in the level of *X* induces fluctuations in *Y* that can cause the cell population to diversify and display two states simultaneously (red scenario) or only one state (black scenario). Noise and the mean level of *X* and the dependency of *Y* on *X* determine whether subpopulations arise or not. **C.** If fluctuations in *X* cause a cell to shift phenotype, then chance events in any of its ancestors might be responsible. This is because some of the molecules in the cell are made in ancestral cells. In the figure, we colour-coded the molecules to mark the ancestor in which they were made. The last two sisters contain molecules from their grandmother. Also note, that cell division can cause variation in molecules between sisters, since the pink-producing cell starts with one molecule more than here sister.

### 2.2 Derivation of the molecule copy number distribution for single cells in terms of their ancestral cell histories

The molecular content of single cells is ever-changing during their cell cycle time. For cells that continuously grow and divide in a non-steady state manner, it is generally impossible to determine the probability distribution of finding a certain number of molecules inside a cell of a given age. However, if the population is in a state of balanced growth, all the probability distributions that characterise this population, such as cell volumes at birth and division, cell ages, generation times and molecule copy numbers, are independent of time (Painter and Marr, 1968). In this growth condition, the number of cells, *N*_*c*_, grows exponentially in time with a constant specific growth rate 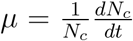. Only in balanced growth, therefore, these distributions are constant in time and may be determined. We assume that populations grow balanced throughout this paper.

The probability that a cell in a population contains *x* molecules of a certain type can be decomposed into two stochastic components. The first component specifies the probability to observe a cell with a certain age *a*, which is described by the statistical theory of cell populations that grow balanced (Collins and Richmond, 1962; Painter and Marr, 1968; Powell, 1964). The second component specifies the probability that a cell at age *a* contains *x* molecules of that type. This probability derives from the theory about the stochasticity of molecular reactions (Van Kampen, 1992; Gillespie, 1992; McQuarrie, 1967). In this work we will merge those two theories.

We denote by *G*_*x*_(*z*) the probability generating function (PGF) of the probability mass function (PMF) of the molecule counts. The PGF determines the probability distribution that a cell drawn at random from a population in balanced growth contains a particular number of molecules of a specific type. *G*_*x*_(*z*) is related to the cell-age distribution *u*(*a*) (a probability density function (PDF)) and *G*_*x|a*_(*z*), the PGF of the probability mass function specifying the probability that a cell contains *x* molecules at age *a* (Schwabe and Bruggeman, 2014a), via

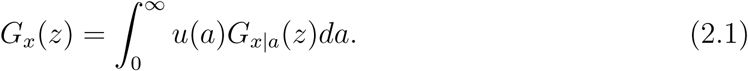

To derive the overall PGF for *G*_*x|a*_(*z*), we first need to consider a single ancestral lineage, including the division times of the mother, grandmother, etc. These PGFs are denoted by 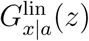. Later we will consider the possibilities of different lineages. When we consider a cell at age *a*, then this cell was born *a* time ago. For the ancestral cells, their ages at division matter most for our theory, i.e., their division times. We shall denote by *T*_1_ the division time of the mother, by *T*_2_ the division time of the grandmother, and so on. The sequence *T*_*i*_ thus marks the sequence of division times of past generations within this specific lineage.

The PGF *G*_*x|a*_(*z*) can itself be expressed as a product of two PGFs. One is associated with the probability of the number of molecules synthesised since cell birth (*a* time ago), and another with the probability that the molecules were inherited from the mother cell, and have ‘survived’ since birth, a period of duration *a*. The latter effect can be considered an example of Lamarckian inheritance. This decomposition is reflected in terms of generating functions as

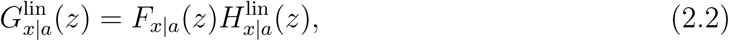

where the PGF 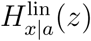 accounts for the probability of the number of molecules obtained from previous generations, and *F*_*x|a*_(*z*) accounts for the probability for the number of newly produced molecules.

The molecules obtained from the mother cell are only degraded in the current cell. Their number at time *a* is described by a Bernoulli distribution with age-dependent parameter *p*(*a*) and PGF 1 – *p*(*a*) + *p*(*a*)*z*, reflecting the probability that a molecule has not yet been degraded at time *a* after birth. This leads to the relation

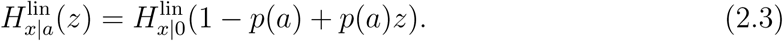

At age 0, the probability for zero synthesised molecules is 1, i.e., *F*_*x|*0_(*z*) = 1. Hence,

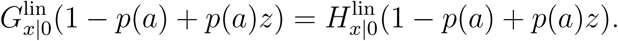

Due to balanced growth, we know that the 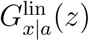 is constant in time: mother and daughter cells have the same distribution. We can therefore relate 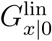 to the molecule count in the mother cell at the time of division *T*_1_, by keeping track which molecules from the mother end up in which daughter. Assuming that for a particular division this probability is equal to a constant *q* for each individual molecule, we have

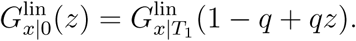

Combined with (2.3), we have

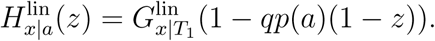

Here we recognise *qp*(*a*) as the probability of a molecule to be partitioned into one of the daughter cells and to survive up to age *a*. Thus, we obtain the following expression

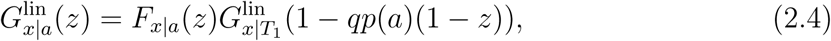

which relates the distribution of molecule counts in the current generation to that of the previous generation. The PGF for the previous generation, 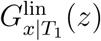, itself satisfies

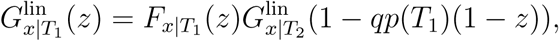

so that

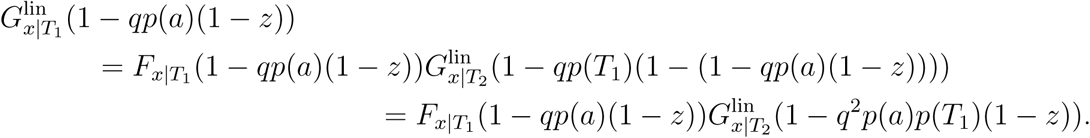

Therefore, equation (2.4) becomes

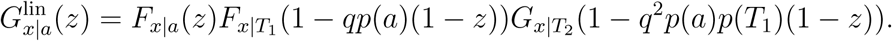

and going back *N* generations leads to the following relation for a single lineage,

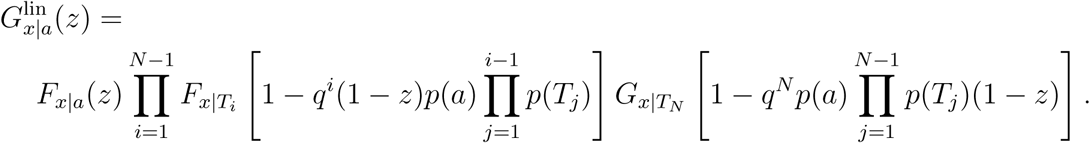

(The terms in large brackets are the arguments of the PGFs, not product terms. The products have a meaningful interpretation for *N ≥* 2, *i ≥* 2; for *N* = 1 and *i* = 1 they should be read as yielding factors of 1.)

As a last step, we need to consider that a single cell may have many possible lineages, each one specified by the division times of its ancestors, *T*_1_, *T*_2_, *…*. Let *f*_*τ*_ (*T)* denote the generation time distribution for the average lineage (Thomas, 2019; Wakamoto et al., 2012). A particular sequence of generation times, i.e., the specific lineage, comes with a probability equal to 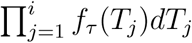. The overall probability that a cell contains *x* molecules at age *a* can now be calculated by integrating over all lineages,

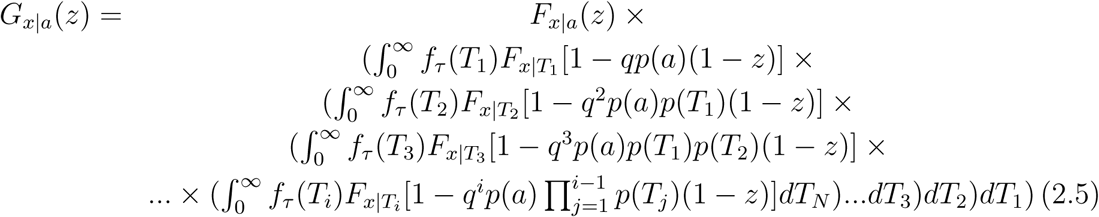

which may be simplified somewhat to

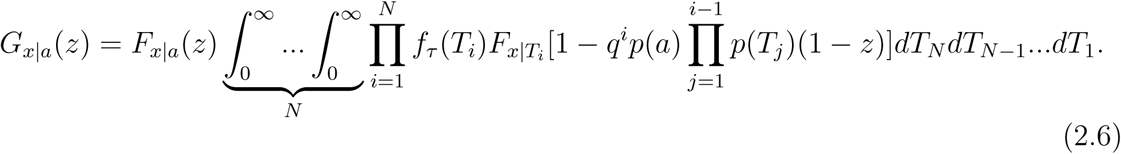

Equation (2.6) is the main equation for this paper. Increasing the number of ancestral generations *N* should improve the approximation of the probability distribution of *x|a*. As we will see, for molecules that have a short “memory” (they are short-lived) considering even one or two generations should give acceptable accuracies. Fluctuations in such molecules may give rise to fate decisions in cells that are only a one or two generations later in the lineage. For long-lived molecules, explaining deviating values requires looking back multiple generations ago.

### 2.3 The rate of molecular memory loss by cell division and molecule degradation

To obtain a good intuition for how many generations we should look back in time to understand the historical events that have shaped the molecular composition of a cell, we derive an expression for the rate of “memory loss”. To do this, we choose 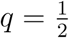, so each molecule has equal probability to be partitioned into either of the two daughter cells. We also choose a specific function for the probability that a synthesised molecule survives (is not degraded) for a period of length *t*, 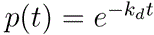, where *k*_*d*_ is a first-order degradation rate constant.

With these two model ingredients, the loss rate can be understood from (2.6) from the term expressing the fraction of molecules that have survived from an ancestor *i* generations ago,

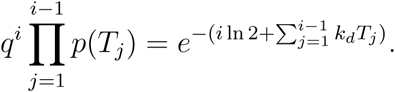

When *i* equals a large value *N* we obtain

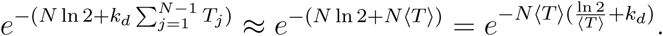

A corresponding macroscopic description of changes in the concentration of this molecule due to dilution by growth and degradation is given by 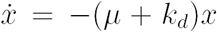 solution 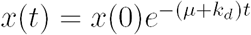. We define *t* := *N* ⟨*T* ⟩ so that,

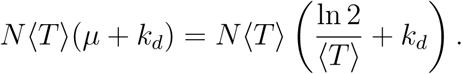

Therefore, an estimate of the average loss rate per generation equals

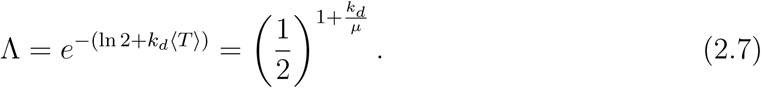

and the fraction of molecules from *N* generations ago hence equals Λ^*N*^. Thus, the dimensionless parameter *k*_*d*_*/µ*, which is proportional to the number of cellular generation times per molecule life time, determines how many generation times one should look back in order to explain the molecule copy numbers that a particular cell contains; the probability that a cell contains molecules that are older than that time becomes negligibly small. Short-lived molecules have a high *k*_*d*_*/µ* value so their memory decays quickly while non-degraded, ‘stable’ molecules have the longest memory since *k*_*d*_*/µ* = 0.

We note that Equation (2.7) cannot be directly substituted in (2.6), because it only applies when we look at histories that are so long that their average generation times matches the mean generation time of the entire population.

### 2.4 The benchmark case of stable molecules

#### 2.4.1 The main equation, (2.6), simplifies for stable molecules

Completely stable molecules, which are never degraded, have the strongest possible lineage effect on copy numbers of an extant cell. This extreme case therefore may serve as a benchmark, against which more realistic scenarios may be compared. The case of stable molecules can be solved explicitly for reasonable assumptions on the mechanism involving molecule synthesis and the generation time distribution *f*_*τ*_ (*T)*, as we will see.

We start from (2.6). If the molecules are not degraded, so that *p*(*a*) = *p*(*T*_*j*_) = 1, we obtain, with *N → ∞*, that

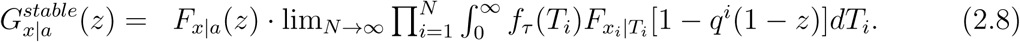

This equation is easier to analyse than (2.6).

#### 2.4.2 The Poisson model for protein synthesis

To gain further intuition, we start by considering the simplest circuit. Let us assume that a molecule is synthesized by a zero-order process, depicted by 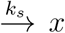. In this case, the probability mass function for the number of molecules synthesized since birth follows a Poisson distribution with parameter *k*_*s*_*a*. Its probability generating function is described by 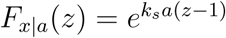, so that

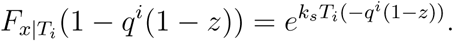

Moreover, we assume the distribution of generation times follows an Erlang distribution,

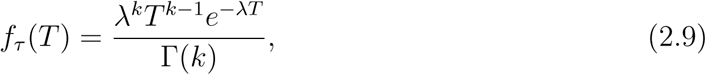

with mean generation time 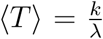 and variance 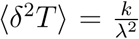, in line with experimental findings (Heerden et al., 2017). A random variable is Erlang distributed with parameters *k* and *λ* if it is the sum of *k* exponentially distributed variables, each with parameter *λ*.

With these two modelling ingredients specified, (2.8) can be explicitly calculated,

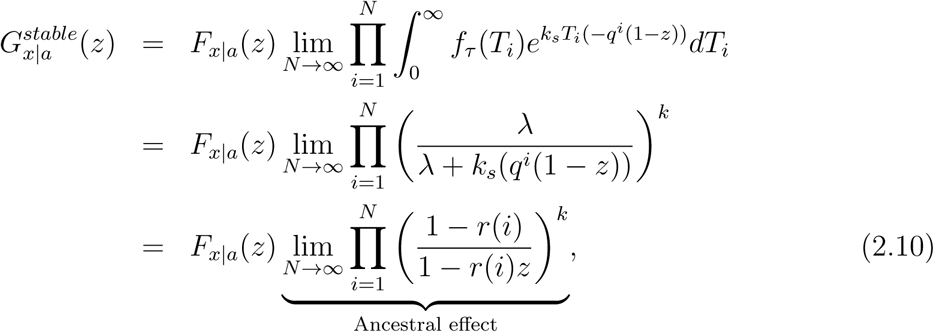

where

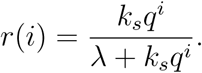

We note that the term 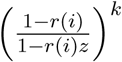 in (2.10) is the PGF of a negative binomial distri-bution with parameters *k* and *r*(*i*), indicating that in this model the number of molecules in a cell equals the sum of a Poisson-distributed number of molecules made in the current generation and of negative-binomially distributed molecule numbers made by ancestral cells.

Thus the number of molecules, *x*_*i*_, in a cell that originate from an ancestral cell *i* generations ago is distributed according to a negative binomial distribution with *i* influencing one of its parameters. It has mean 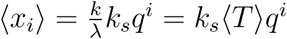, which equals the multiplication of the mean number of molecules made in this cell, *k*_*s*_⟨*T*⟩, by the total dilution due to cell divisions, i.e., *q*^*i*^. Hence, in this way we have formally established that for completely stable molecules, on average a cell that is about to divide contains molecules that are for 50% made by itself, 25% by its mother, 12.5% by its grandmother, etc (as one would have expected, of course).

For a negative binomial distribution with parameters *k* and *r*(*i*), the variance equals the mean divided by 1 – *r*(*i*). Hence, here 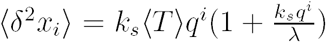, with *k*_*s*_*q*^*i*^*/λ* as the number of surviving molecules that were made in time interval 1*/λ*. Adding to this that we have 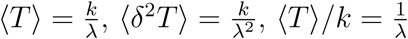, we get

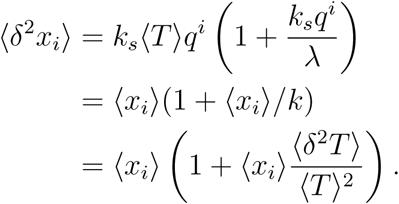

Therefore, the noise in that number of molecules equals

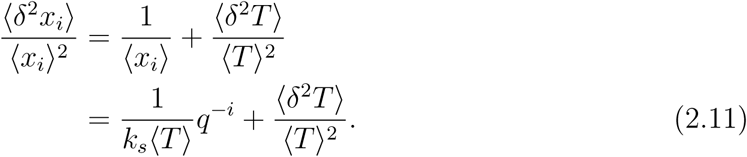

This is an important relation for our intuition. It indicates that: 1. the noise in the generation time distribution codetermines the heterogeneity in the copy numbers; 2. the noise is decreased if cells produce more molecules during single cell cycles, but not to zero; 3. the noise increases with the number of past generations (*q*^*-i*^ increases with *i*). The last conclusion implies that we lose certainty – ‘information’ – when we consider older chance molecular events; they are expressed as a longer product of probabilities. Note however, that while the noise increases, its effect on the current cell could still decrease because the number of molecules decreases.

We show how the ancestral history disappears in a given cell as function of the number of past generations in Figure 2. Figure 2A shows the probability distribution of molecule copy numbers in the entire population of growing cells. In Figure 2B, the probabilities are shown that cells from the same population contain a certain number of molecules from ancestral cells *i* generations back. The probabilities of retaining molecules from past generations decreases.

**Figure 2:**
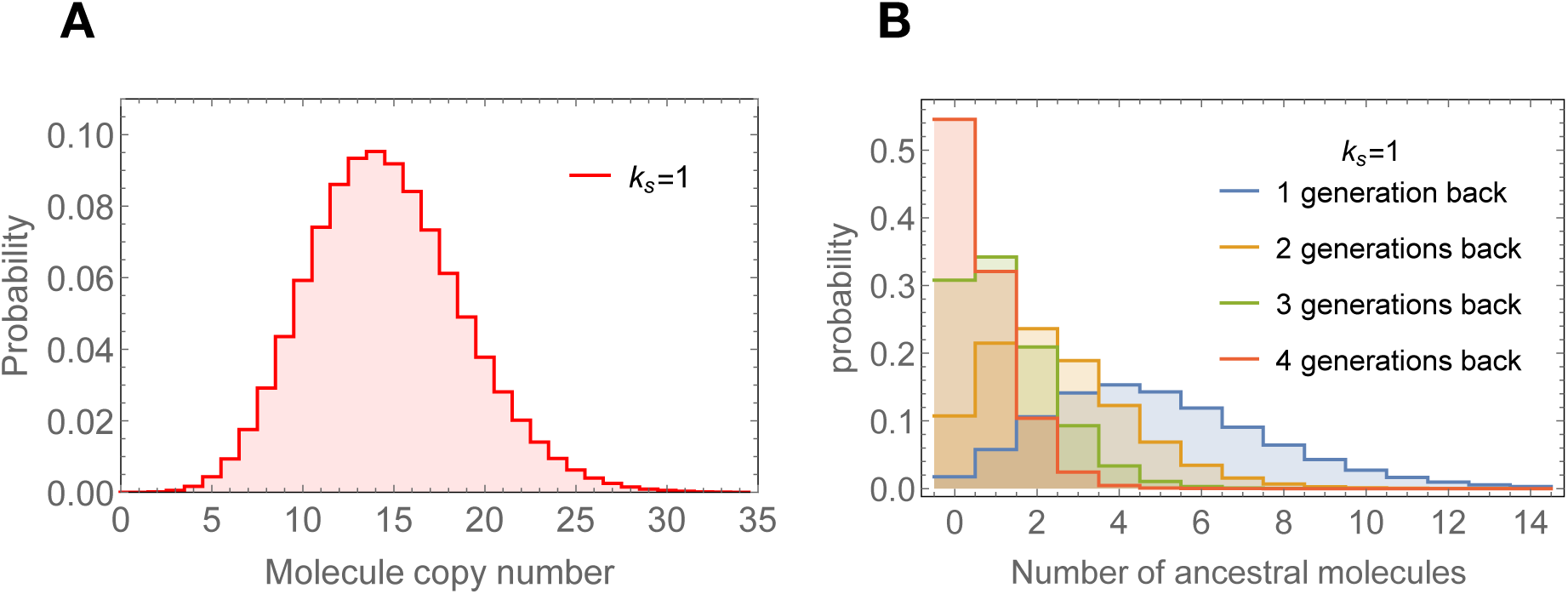
Illustration of how the molecule copy number distribution of a cell can be viewed as a sum of contributions made by its ancestral cells. **A.** The molecule copy-number distribution according to (2.10) with parameters *k*_*s*_ = 1 *min*^−1^, *k* = 10, *λ* = 1 *min*^−1^, and 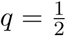. This models a cell with a realistic generation times (mean 60 *min* and coefficient of variation of 30 %) and a low copy number of proteins, e.g. transcription factors. Cases of high protein expression are considered below. Calculation were done at the mean cell age. **B.** Contributions of ancestral cells to the molecule copy-number distribution shown in Figure A, according to (2.10). Calculations were done at the mean cell age.

#### 2.4.3 The macroscopic description of molecule copy numbers can be a poor approximation of the stochastic description

Using relation (2.10), we can determine the mean and variance of the copy number of molecules. The mean of a sum of random variables equals the sum of the means; therefore, the mean copy number at a particular cell age equals,

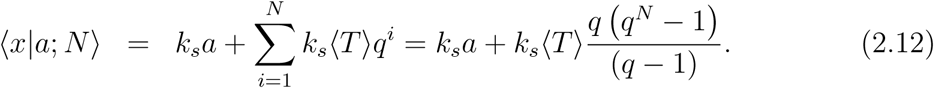

For *q* = 1*/*2, we have that *q*(*q*^*N*^ – 1)*/*(*q* – 1) *→* 1 for *N → ∞*, so that

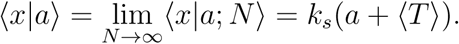

The term *k*_*s*_⟨*T*⟩ equals the number of molecules at cell birth and *k*_*s*_*a* equals the number of molecules made since birth.

The copy-number variance at a particular age, ⟨*δ*^2^*x|a*⟩, equals the sum of the variances of the molecules made in different cell generations since those random events are independent in our model. Since we are considering a zero-order synthesis process, the number of molecules made in the cell follows a Poisson process and therefore has variance equal to the mean, ⟨*δ*^2^*x|a*⟩ = ⟨*x|a*⟩ = *k*_*s*_*a* and the variance of the ancestral molecules that survived *i* generations, equals *k*_*s*_⟨*δ*^2^*T* ⟩*q*^*i*^ (*k*_*s*_*q*^*i*^ + *λ*). Therefore,

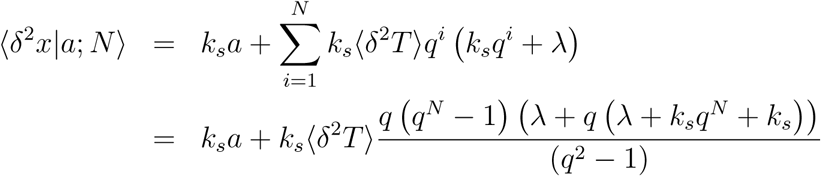

so that for *q* = 1*/*2,

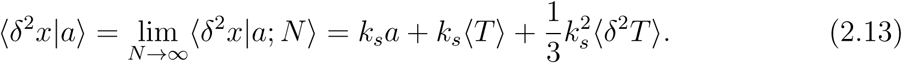

We can now establish the overall mean and variance by integration over the age distribution. For the mean we find

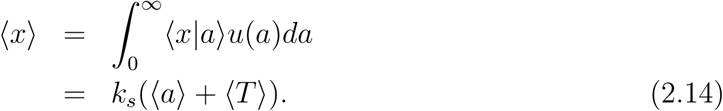

The variance needs to be decomposed into

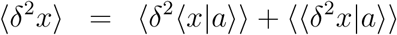

where

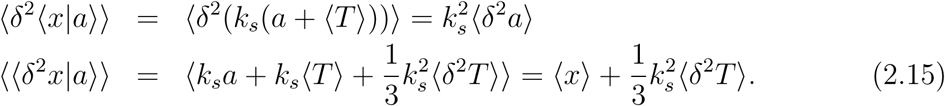

Thus we still need to calculate the mean ⟨*a*⟩ and the variance ⟨*δ*^2^*a*⟩ of the cell age to be able to calculate the quantities mentioned above.

Following (Painter and Marr, 1968), we can calculate the population growth rate *µ* from 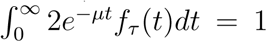, after which we can determine the age distribution from 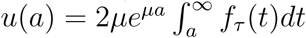 For the specific choice of an Erlang distribution for *f* _τ_(*T)* chosen earlier, see (2.9), the growth rate equals 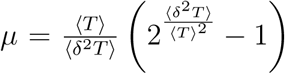. This indeed equals the macroscopic estimate 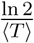 when ⟨*δ*^2^*T*⟩*→* 0. Now that we have found *µ*, we can determine

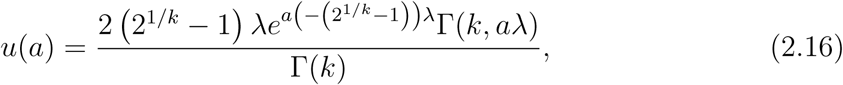

which has as mean value

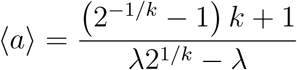

and variance

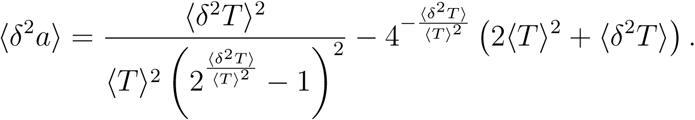

Using these identities, the values for the mean and variance of the copy number of *X* (2.15) become

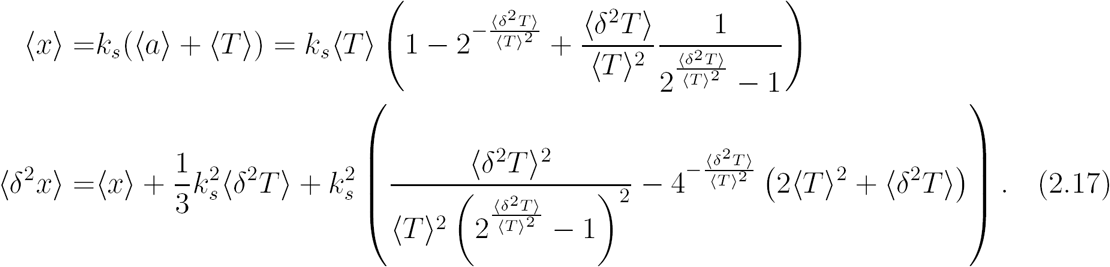

When we take the limit ⟨*δ*^2^*T* ⟩ *→* 0 then 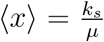 and ⟨*δ*^2^*x*⟩ = ⟨*x*⟩ + ⟨*x*⟩^2^(1 − 2(ln 2)^2^) *≈* ⟨*x*⟩ + 0.039⟨*x*⟩^2^. The mean value agrees with the deterministic model 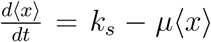 for constant synthesis rate *k*_*s*_ and dilution by growth, which indeed has as steady state 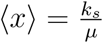, indicating that the macroscopic approximation assumes ⟨*δ*^2^*T*⟩*→* 0.

When this macroscopic ODE system is modelled probabilistically with chemical master equations and then linearised, using a method called linear noise approximation (LNA; see (Van Kampen, 2003)), the mean and variance around this steady state may be determined analytically, showing that they are equal, ⟨*x*⟩ = ⟨*δ*^2^*x*⟩ (Elf and Ehrenberg, 2003; Bruggeman et al., 2009). Thus, we conclude that variance estimation with LNA misses the term ⟨*x*⟩ ^2^(1 − 2(ln 2)^2^) when ⟨*δ*^2^*T*⟩ *→*0 because cell growth is not modelled in sufficient detail. Therefore, copy-number noise estimations of biochemical circuits in single cells that are members of cell populations that grow balanced, using LNA, lack the term (1 − 2(ln 2)^2^). This amounts to 20% coefficient of variation in molecule copy number and can exceed the value 1*/*⟨*x*⟩ by orders of magnitude.

Equation (2.17) also indicates that the variance in molecule copy numbers is increased due to heterogeneity in generation times. (This is because cells having a deviating generation time have had deviating times for molecule synthesis and therefore made different amounts.) The increased variance in copy numbers, due to generation time variation, indicates that *molecular* non-phenotypic heterogeneity is enhanced by the stochasticity of *cellular* processes.

### 2.5 Noise in molecule copy numbers always exceeds a noise floor that is set by the probability distribution of generation times

The equations for the mean and variance in copy numbers, (2.14)–(2.15), indicate the existence of a ‘noise floor’, the lowest possible noise value, for any generation time distri-bution,

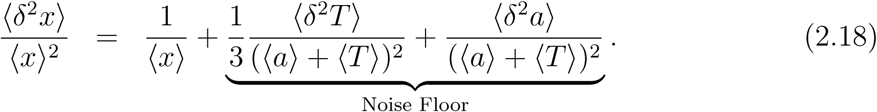

This result indicates that the noise floor, which is also found in experimental data (Wolf et al., 2015; Taniguchi et al., 2010), cannot be explained from biochemical reaction kinetics alone: the stochasticity associated with cell growth is responsible for its existence. Its theoretical value is actually quite close to experimental data (Kempe et al., 2015). We have also seen that, in the case of an Erlang-distributed generation time, when the variance of the generation time becomes zero, the second term in the noise floor equals 1 − 2(ln 2)^2^.

Thus, noise in copy numbers cannot become zero in asynchronously growing cell populations. The second term of the noise floor arises because molecule copy numbers increase with cell age, and a mixture of cell ages is always found in such conditions, so that heterogeneity in copy number immediately follows. Note that molecular noise considered in terms of concentrations does not have this noise term (Schwabe and Bruggeman, 2014a). The first term of the noise floor, however, is due to variation in cell cycle times and will persist even if concentrations are considered.

The noise floor argues against an often used rule of thumb, i.e 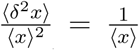, that suggests that copy number noise in abundant molecules is negligible. The existence of the noise floor implies that the origins of fate decisions can come from deviating copy numbers of abundant molecules. This can underlie cell-fate decisions leading to persister cells in bacterial cultures (since those have been attributed to stochastic events associated with metabolism (Maisonneuve et al., 2013)) provided that those phenomena are sensitive to copy number noise and not to concentration noise, as concentration noise does not appear to have a large noise floor (Schwabe and Bruggeman, 2014a).

### 2.6 How do random bursts of translational activity in ancestral cells influence cell fate decisions?

Transcription and translation bursts are a potent mechanism to enhance molecular heterogeneity across isogenic cells (Raj and van Oudenaarden, 2008). Here we use the classical model for translation bursts (Shahrezaei and Swain, 2008; Dobrzynski and Bruggeman, 2009) in the context of ancestral history effects on the randomness of molecule copy numbers in single cells obeying balanced growth statistics.

The model describes a gene switching between an OFF and an ON state. Only during the ON state transcription occurs. Due to the switching of the gene, transcription activity will switch at random times from ON to OFF, and back, leading to inactive and active periods of random durations. During the active periods, synthesis and degradation of mRNA and proteins occurs while during the inactive period mRNA synthesis stops.

We shall be considering the following molecular circuit, with mRNA and protein having copy numbers *n*_*m*_ and *n*_*p*_, respectively, and assume it to be embedded in a single cell that is a member of an isogenic population of cells that grows balanced,

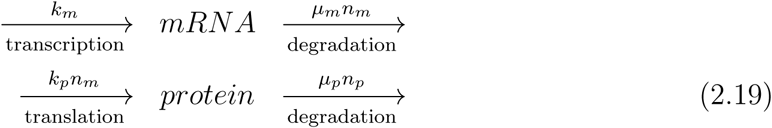

#### 2.6.1 Stochasticity of translation bursts

A translation burst size is defined as the number of proteins made during the life time of a *single* mRNA. Thus, we are considering (note that *n*_*m*_ = 1),

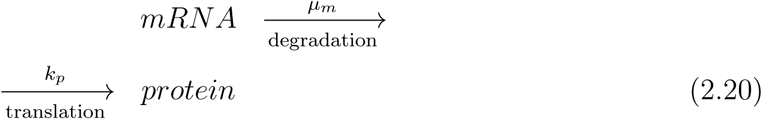

Since the times between consecutive protein production events are exponentially distributed, with mean time 1*/k*_*p*_, the number of proteins made in a time interval of length *t* follows a Poisson distribution (Schwabe et al., 2012) with mean and variance *k*_*p*_*t*; hence, 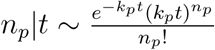. The duration of the protein production time is random itself and equals the life time of the mRNA, which we assume to follow an exponential distribution with mean *µ*_*m*_. The resulting probability distribution of the burst size, *b* therefore equals

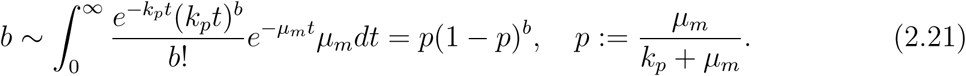

Thus, the burst size follows a geometric distribution with mean

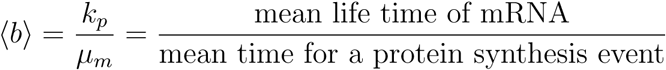

and variance ⟨*δ*^2^*b*⟩ = ⟨*b*⟩^2^ + ⟨*b*⟩. We note that translation bursting causes large heterogeneity in protein copy numbers per cell because its noise equals 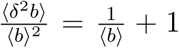, which should be compared to 0 when bursts occur of a fixed size.

A useful relation for later is that the PGF for a geometric distribution equals 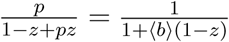 and that a sum of *N* independent geometrically distributed random variables equals a negative binomial distribution, such that its PGF equals 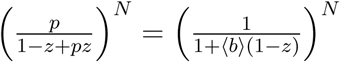.

#### 2.6.2 Probability distribution of molecule copy numbers

Following (Shahrezaei and Swain, 2008) the PGF for the number of proteins made by the circuit in (2.19) during a time interval of length *t* is given by

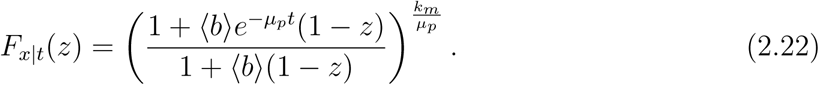

If the cell had *n*_0_ molecules to start with then this PGF should be multiplied with the ‘survival’ term 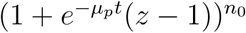.

If the proteins are not degraded then the previous PGF (2.22) simplifies to

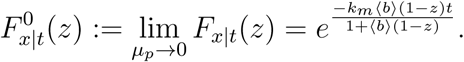

Under this assumption the distribution of molecule copy numbers in a growing cell population can be obtained from (2.8). We first calculate the integral,

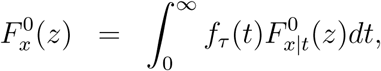

which, using the choice *f*_*τ*_ (*t*) ∼ *Erlang*(*k, λ*) becomes

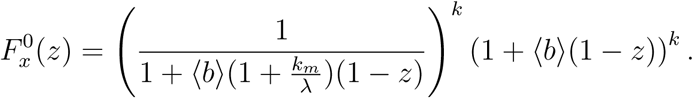

We note that this PGF equals the ratio of the PGFs of two negative binomial distributions, but we do not understand fully the physical interpretation of this insight. To determine the probability generating function of the molecule copy numbers in a growing population of cells we would have to calculate 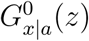. The obtain the noise floor, however, we only need to derive an expression for the mean and variance of this molecule copy number.

We start by deriving those quantities conditional on the age. Since the distribution of *x*_0_|*a* has PGF 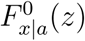 and that of *x*_*i*_ is 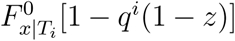, using reasoning identical to that for the Poisson model in Section 2.4.3,

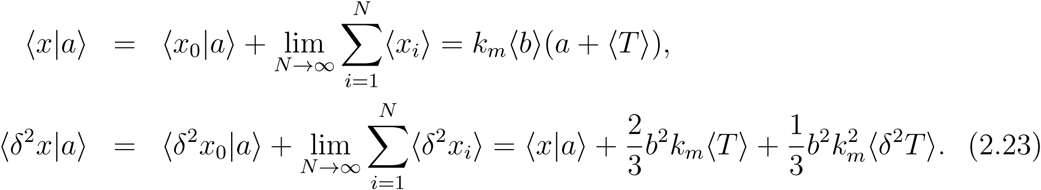

For the mean and variance we obtain,

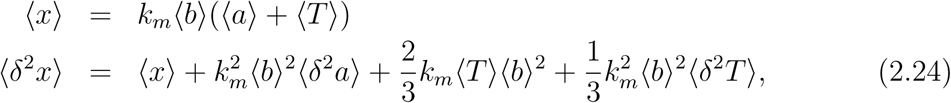

which are completely general and independent of the mathematical definition of the interdivision time distribution. The noise in molecule numbers is therefore given by the following expression,

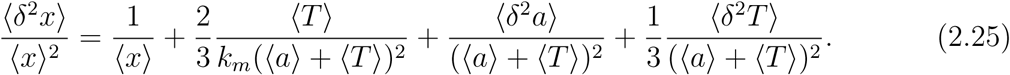

To derive the noise floor we have to consider the noise at ⟨*x*⟩ → ∞. If we achieve this by changing the transcription rate *k*_*m*_, as one would normally do, then the last two terms in the previous equation remain,

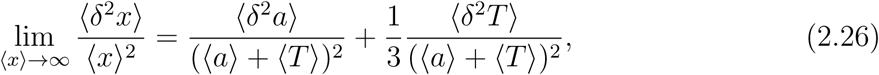

which is identical to the noise floor (2.18) that we found earlier for a model that did not include transcriptional bursts. The difference between these models is that at low and moderate levels copy numbers the translation bursts model generates more noise.

Thus, we find again that the noise floor is completely determined by the stochasticity associated with cell growth and that therefore cell fate decisions may also depend on molecules with high copy numbers, for which 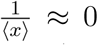. (Provided that those decisions depend on copy numbers and not on concentrations.)

Figure 3 summarises the main insights from the analysis of noise floors (eq. 2.25). In Figure 3A the noise floor is shown as function of the mean copy number, which ranges from a few per cell, for transcription, to hundreds for signalling proteins and (tens of) thousands for metabolic proteins. Thus at least about 20% of phenotypic variation is found for molecule copy numbers in growing cell populations that is entirely due to those populations consisting of cells having different sizes, i.e., ages, and that from birth to division the number of molecules doubles, during balanced growth. Figure 3B shows the dependency of the noise floor on the coefficient of variation of the generation time distribution *f*_*τ*_ (*t*). Large variation in generation times contributes to molecule copy number heterogeneity because cells that live longer make more molecules than cells living shorter.

**Figure 3:**
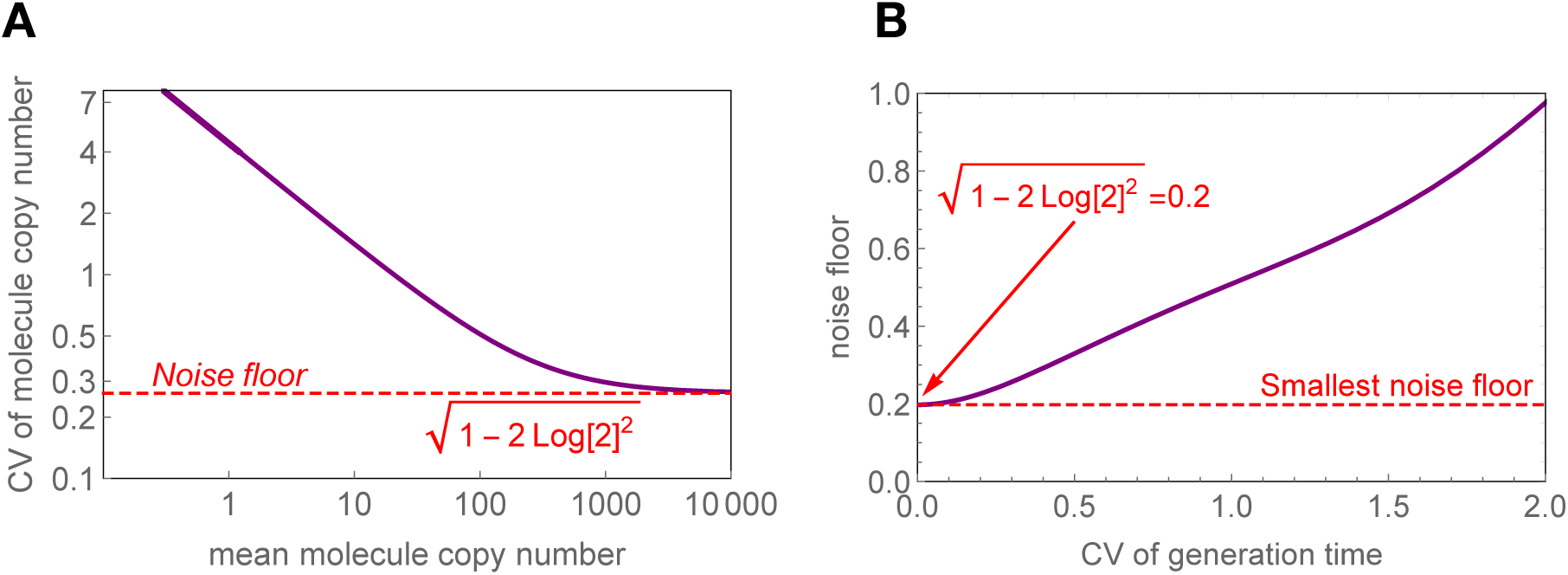
Noise floor of molecule copy-numbers. **A.** Calculation of the relation between the coefficient of variation and the mean of the molecule copy-numbers. This relation was calculated by varying the growth rate; alternatively we could have changed the rate of mRNA synthesis. The other parameters were to *k*_*s*_ = 2 *mRNA/min, k*_*p*_ = 4 *proteins/min, µ*_*m*_ = 0.1 *min*^−1^. In bacterial cells, transcription factors occur as a few to tens of copies of per cell whereas metabolic proteins occur as hundreds to tens of thousands of copies per cell. **B.** The noise floor is only determined by the coefficient of variation in the cellular generation time, which has experimentally been found to be ≈ 0.3 (e.g. Heerden et al. (2017)). Those two plots were generated using equation 2.25 and by assuming that the interdivision time follows an Erlang distribution.

### 2.7 Influence of chance events associated with cell growth

Growth rate influences many aspects of cell physiology (Klumpp et al., 2009). One of its effects is that it sets the generation time, which limits the time that cells have for molecule synthesis from their birth to division. The analysis leading to (2.25) indicates that if cells would tune mRNA synthesis with growth rate in order to compensate for the reduction in copy numbers of mRNA and protein molecules due to shorter generation times, noise in protein expression would be independent of growth rate. We think it is unlikely that cells can coordinate mRNA and/or protein synthesis with such precision. Molecular noise is therefore expected to be growth rate dependent. For instance, experimental data indicates that noise of a constitutively expressed protein increases with growth rate in *B. subtilis* (Nordholt et al., 2017).

Genes that are turned off by repressors do occasionally make proteins if the repressor falls off its regulatory site and a RNA polymerase binds instead (Choi et al., 2008). This “leaky gene expression” can lead to appreciable protein accumulation at low growth rate, because then generation times are long. This is shown in Figure 4AB. At high growth rate, the probability of a leaky expression event of a repressed gene during a single generation time is small. Few cells will therefore experience such faulty events and the probability for a cell to have zero molecules is then high. At low growth rate, however, leaky gene expression events become more probable during a single generation time and more cells will experience leaky gene expression events occurring purely by chance. These cells accumulate proteins that they do not require now.

**Figure 4:**
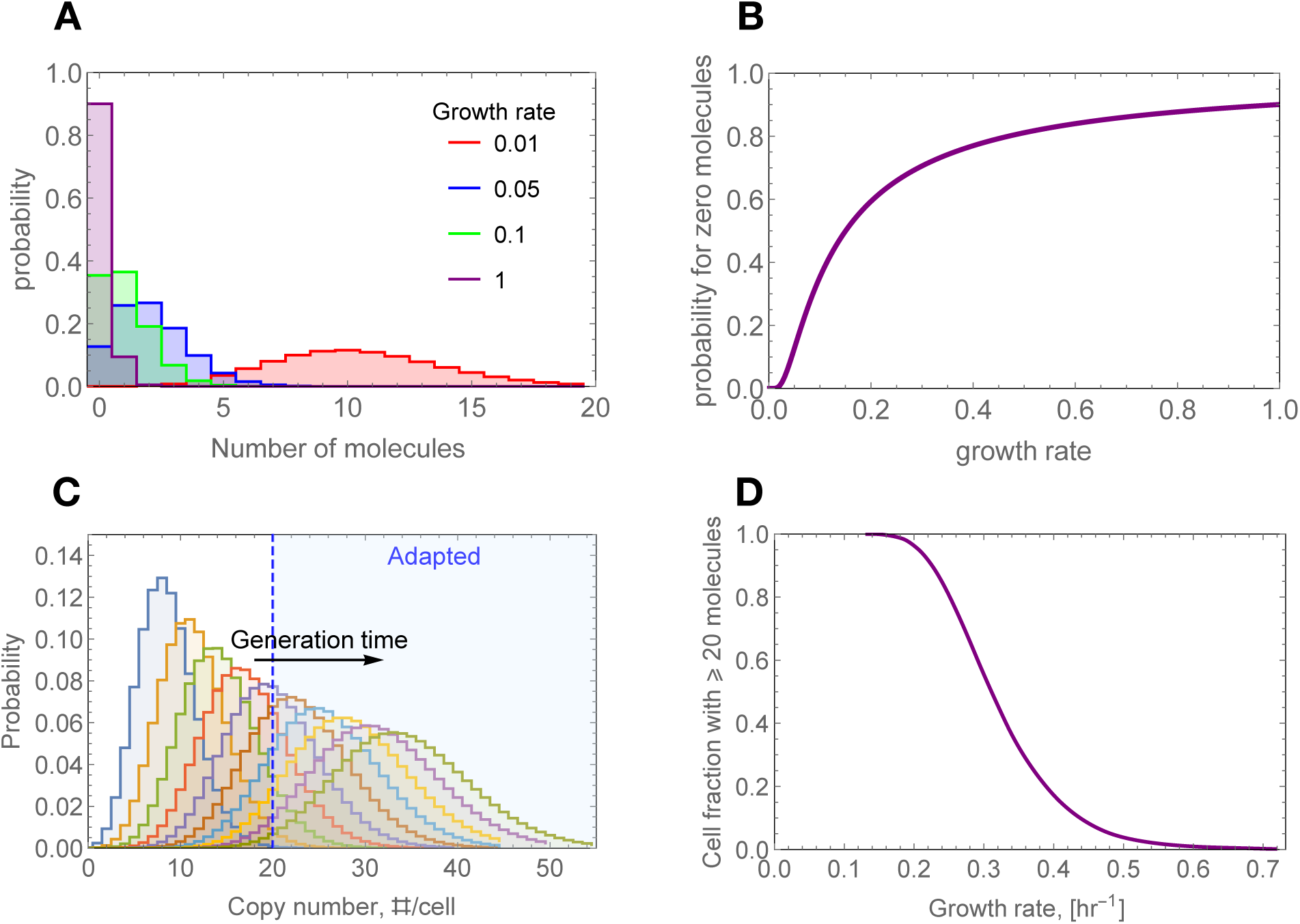
The influence of growth rate on ‘leaky’ gene expression. **A.** The growth rate influences the molecule copy number distribution. Since higher growth rate imply shorter generation times, less molecules are made by cell during the period from birth to division. Here we consider a gene that sometimes makes an mRNA under conditions when it is supposed to be off – it is a ‘leaky’ gene. Calculations were done at the mean cell age, using (2.10). **B.** At low growth rate single cells accumulate protein via leaky gene expression while this effect is negligible at high growth rate. Cell physiology can therefore be expected to be more noisy at low growth rate. Calculations were done at the mean cell age.

This accumulation of (apparently) unnecessary proteins due to leaky gene expression at low growth rate, can lead to spontaneous adaptation of cells, because cells could accumulate proteins above a threshold level. In this way, a contribution to adaptation to a new future condition appears purely by chance. This is illustrated in Figure 4CD. The probability distributions of molecule copy numbers were calculated at different growth rates. The probability that a cell contains more than 20 molecules, which we set to be the threshold fraction for adaptation, decreases with growth rate. This indicates that some cells have a spontaneous adaptation capacity at low growth rate, due to chance events (Choi et al., 2008), and that this cell fraction decreases with growth rate.

## 3 Discussion

Since mother cells pass on average 50% of their molecular content on to each daughter cell, the inherent randomness of molecular and cellular processes of one cell can affect the fate of its progeny cells. The lineage history of a cell therefore matters when we aim to understand why it displays a qualitatively different phenotype than its kin cells. This ‘Lamarckian inheritance’ mechanism plays an important role in the emergence and maintenance of phenotypic heterogeneity in growing cell cultures. In this paper, we related the molecule content of a cell to that of its ancestors.

We combined stochasticity at the molecular level with stochastic cellular processes. In this way we obtained a full probabilistic description of molecule copy numbers in exponentially growing cells. The condition of balanced growth is a requirement for our theory, because only under those conditions all the probability distributions of intensive characteristics of the growing cell population are time-invariant. The theory relates the current molecular state of the cell to that of its ancestors along its lineage, taking into account molecule turnover and partitioning during cell division. Our main equation (2.6) is completely general and can be used with any stochastic biochemical reaction network. It will not always be analytically solvable, in that case numerical simulations will be required.

The theory considers the statistics of molecular and cellular events along lineages. The generation times of cells play an important role. First, whether a single ancestral molecule survives in offspring cells depends on whether it is degraded before it is partitioned into the daughter cells. Therefore its survival depends on the generation time of the mother. Second, since the generation time sets the total synthesis time for cells to make molecules, cells with long generation times generally make more molecules. Thus, the variation of generation times between cells plays an important role in our theory. If those generation times show great variation then the population will be necessarily more heterogeneous too. We have little molecular understanding at the moment of what sets the generation time of a cell and how its variance is determined.

Our theoretical results mostly concern the case of infinitely stable molecules that do not degrade, but only dilute by cell division. The effect of ancestral stochasticity is greatest for such stable molecules, of which metabolic proteins are good examples.

The theory indicates that variances in copy number and in generation times matter to molecular heterogeneity. It has been known for a long time that nonlinear kinetics gives rise to approximative ordinary differential equation (ODE) models because of the lack of moment closure, i.e., the ODEs for the mean copy number also depend on the variance of the copy number (Goel and Richter-Dyn, 1974). Here, we have shown that deterministic ODE models of the dynamics of molecular circuits inside living cells fail even for the simplest possible zero-order reaction kinetics. Also the variance of the generation time matters: the mean copy number only equals the macroscopic ODE result when the variance of the generation times equals zero.

We found a noise floor that specifies a lower bound to the noise in highly abundant molecules, with copy numbers exceeding a 100 or so. Since 1 molecule per bacterial cell, such as *E. coli*, is approximately equal to a concentration of 1 nM, our result indicate that the noise floor applies to molecules that have concentrations in the *µ*M range or higher. Those are mostly metabolic proteins and low-molecular weight molecules, e.g., intermediates of metabolism. Transcription factors and signalling proteins are generally lower in concentrations and shall experience higher noise levels than the noise floor.

The noise floor arises for two reasons. Firstly, in a cell population at balanced growth extant cells have different ages: some are just born, others are halfway through their cell cycle and others are about to divide. Since the copy number of each molecule is doubled on average during a single cell cycle, the molecule content of cells increases with their age. Heterogeneity in copy number therefore arises because of the heterogeneity in cell ages. This effect is independent of the mean copy number of a molecule so it persists when the mean copy number approaches infinity. Secondly, the variation in generation times causes variation in the number of molecules made by cells during their cell cycle. We note that the first effect also arises when all cells have the same generation time (Schwabe and Bruggeman, 2014b). Cell ages also have generally greater variance when the generation times are more variable amongst cells.

Copy number variation is higher for cell cultures with longer generation times. The theory therefore indicates a possible mechanism to induce heterogeneity, thereby increasing the adaptive capacities of microbial cell populations. It indicates how leaky gene expression of a repressed gene can lead to a bet-hedging type of adaptive behaviour at low growth rates that diminishes at high growth rate. The relationship between growth rate and protein copy number heterogeneity is important for microbiology. It appears to be associated with the adaptation of microorganisms to new nutrient conditions (Choi et al., 2008; Boulineau et al., 2013; van Heerden et al., 2014; Solopova et al., 2014; Kotte et al., 2014) and likely also the formation of persister cells (Balaban et al., 2004; Harms et al., 2016) as critical ppGpp fluctuations for persister cell formation are likely more probable at low growth rate when the mean ppGpp concentration is higher.

From this work we draw several conclusions. Fluctuations in ancestral cells that led to deviating phenotypes can propagate along a lineage tree of growing cells, causing the formation of a subpopulation of cells, because mother cells pass on part of their molecular state to their daughters. The variation of generation times between cells plays an important role in copy-number heterogeneity in growing cell populations; it contributes to a copy-number noise floor that exists for each molecule. Finally, Lamarckian inheritance appears to play an important role in setting the heterogeneity and adaptive capacities of growing cell populations.

## 4 Acknowledgements

Daan de Groot was supported by the NWO VICI grant 865.14.005.

